# Tracking canopy gap dynamics across four sites in the Brazilian Amazon

**DOI:** 10.1101/2022.09.03.506473

**Authors:** Eric Bastos Gorgens, Michael Keller, Toby D Jackson, Daniel Magnabosco Marra, Cristiano Rodrigues Reis, Danilo Roberti Alves de Almeida, David A. Coomes, Jean Pierre Ometto

## Abstract

**Background information to give context to the study:** Canopy gaps are the most evident manifestation of how disturbances disrupt forest landscapes. The size distribution and return frequency of gaps, and subsequent recovery processes, determine whether the old-growth state can be reached.

**The aim or research question:** We used remote sensing metrics to compare the disturbance regime of four Amazon regions based on the size distribution of gaps, their dynamics and geometric characteristics.

**A brief summary of the methodology used:** We assessed gap dynamics at four sites in the central, central eastern, southeastern, and northeastern regions of the Brazilian Amazon using repeated airborne laser scanning surveys. We developed a novel analysis to quantify four possible stages of gap dynamics: formation, expansion, persisting and recovering. For that, we overlapped layers of gap locations from two consecutive airborne laser scanning surveys.

**Key results with some significance measures:** The gap fraction in our study sites varied between 1.26% to 7.84%. All the sites have similar proportion of gaps among size classes. What notably changed between sites was not the gap size-distribution, but the relative importance of stages of gap dynamics. Growing and persisting rates were greatest in the site with the stronger seasonal variation in climate, lower annual precipitation, higher mean wind speed and higher solar radiation.

**The conclusions, which address the main aims:** The concept of stability reflects the tendency of a system to quickly return to a position of equilibrium when disturbed. We showed that gap dynamics varied among sites, with one example of low recovery rate contrasted to three other sites with faster recovery. Our results support that such as assessing the size distribution of gaps, investigating their return frequency and severity is crucial for understanding forest dynamics at the landscape and regional scales.

## Introduction

Gaps are a manifestation of disturbance in forest landscapes (Whitmore 1989, Muscolo et al 2014, Marra et al 2018). We can understand disturbance as “any relatively discrete event in time that disrupts ecosystem, community or population structure and changes resources, substrate availability, or the physical environment” (White and Pickett 1985). Disturbance creates a landscape with patches of different successional stages. The size distribution and return frequency of disturbance events, and subsequent recovery processes, determine to a large extent the spatial scale over which an old-growth steady state develops (Chamber et al 2013).

Gaps can be formed by endogenous senescence of single canopy trees or branches (Hartshorn 1978, Schaetzl et al 1988, Arellano et al 2021). Recent studies, often using remote sensing, demonstrate that exogenous disturbance agents can also cause or amplify gaps. For example, wind can snap or uproot trees (Espirito-Santo, et al. 2010; Negrón-Juárez et al 2011, Toledo et al 2013, Marra et al 2014; Silverio et al. 2018); lightning strikes can cause tree mortality and branch fall (Gora et al 2020); and the frequency of extreme rainfall events may increase the frequency of gaps (Peixoto 2019, Araújo et al 2021, Cushman et al 2022).

Hartshorn (1978) estimated that 75% of tree species in tropical forests depend on canopy gaps for successful regeneration, making gap dynamics a fundamental process in tropical forest ecology. A forest stand is in constant flux, with long stretches of biomass accumulation often punctuated by episodic disturbances, cautioning us not to assume that forests are in steady-state (Chambers et al 2013). Pioneering studies demonstrated that the frequency of gaps in old-growth tropical forests ranges from 0.5 to 2.0% of area per year (Brokaw 1982). Recent studies continue to find results in this range at various locations including 1.3% yr^-1^ in French Guiana, 1.4% yr^-1^ near Manaus, Brazil, (Peixoto 2021), and approximately 2.1% yr^-1^ in Central Panamá (Araújo et al 2021). The latter study conducted over 5 years, revealed that gap formation over a period of five years was highly stochastic, with 23% of the events recorded in a 42-days interval.

Most tropical forest gaps are relatively small and may not provide sufficient light to initiate secondary succession, nor to shift community composition (Hubbell et al, 1999). Succession-inducing events (usually associated with large gaps, >1000 m^2^) are not well represented on traditional forest inventory approaches. It is important to consider landscape- scale processes when studying old-growth forest ecosystems. Chambers et al (2013) suggested plots should be larger than 10 ha to maximize the detection of temporal disturbance trends.

The use of airborne laser scanning to cover large areas with high density of returns and multi-temporal flights provide powerful data to study canopy dynamics. Several studies have characterized gap size frequency in tropical forest using single lidar acquisitions (e.g. Zhang 2008, Kellner & Asner 2009; Asner et al 2013, Espirito-Santo et al. 2014; Hunter et al 2015, Dalagnol et al 2019, Dalagnol et al 2021). A limited number of studies have used multitemporal lidar to map gap creation directly (Hunter et al 2015, Leitold et al 2017). Detailed work in Central Panama has used drone carrying a RGB camera to document canopy dynamics in a 50-ha plot at high temporal frequency (monthly) (Araújo et al 202).

The Sustainable Landscape Brazil Project has repeatedly surveyed Amazonian sites with airborne laser scanning. This data set gave us a unique opportunity to assess gap dynamics across Amazonia (data freely available at: https://www.paisagenslidar.cnptia.embrapa.br/webgis/). Here, we studied the size distribution of gaps and gap dynamics at four Amazonian upland forest sites, including a site where we discovered the tallest trees known from the Amazon region (Gorgens et al 2019). Using airborne lidar collected at two time periods in each of our study sites, we addressed three questions: (Q1) Do the distributions of gap sizes vary among sites and are the differences preserved over time? (Q2) Are the rates of new gap formation, gap expansion, gap persistence, and gap recovery significantly different among sites? (Q3) Are losses of canopy height due to gap formation and expansion compensated by increased growth of survival and successional trees? Our findings give us new insights into gap dynamics at the landscape scale and across a climate and environmental gradient of Amazon.

## Material and Methods

### Study sites

We assessed gap dynamics at four study sites in the Brazilian Amazon: Ducke (1,205 ha), Tapajos (1,026 ha), Tanguro (590 ha), and Jari (813 ha) (Figure 1). All study sites were covered with old-growth forest, with no anthropogenic disturbances detected in the Landsat archive (approximately 40 years). The northeast site (Jari) is located in the Jari basin, between the states of Pará and Amapá states. The southeast site (Tanguro) is located in the state of Mato Grosso, in the Xingu basin (Silverio et al. 2019). This region is marked by the transition between Cerrado and Amazon. The central-east site (Tapajos) is located in the state of Pará, in the Tapajós basin. The fourth site is located in central Amazon (Ducke), in the state of Amazonas at the Amazonas basin. The sites cover a range of climatic conditions, which is detailed in Table 1. For example, annual average precipitation ranges from about 1,600 mm year^-1^ in Tanguro to ~2,300 mm year^-1^ at Jari (Table 1).

**Figure 1.**
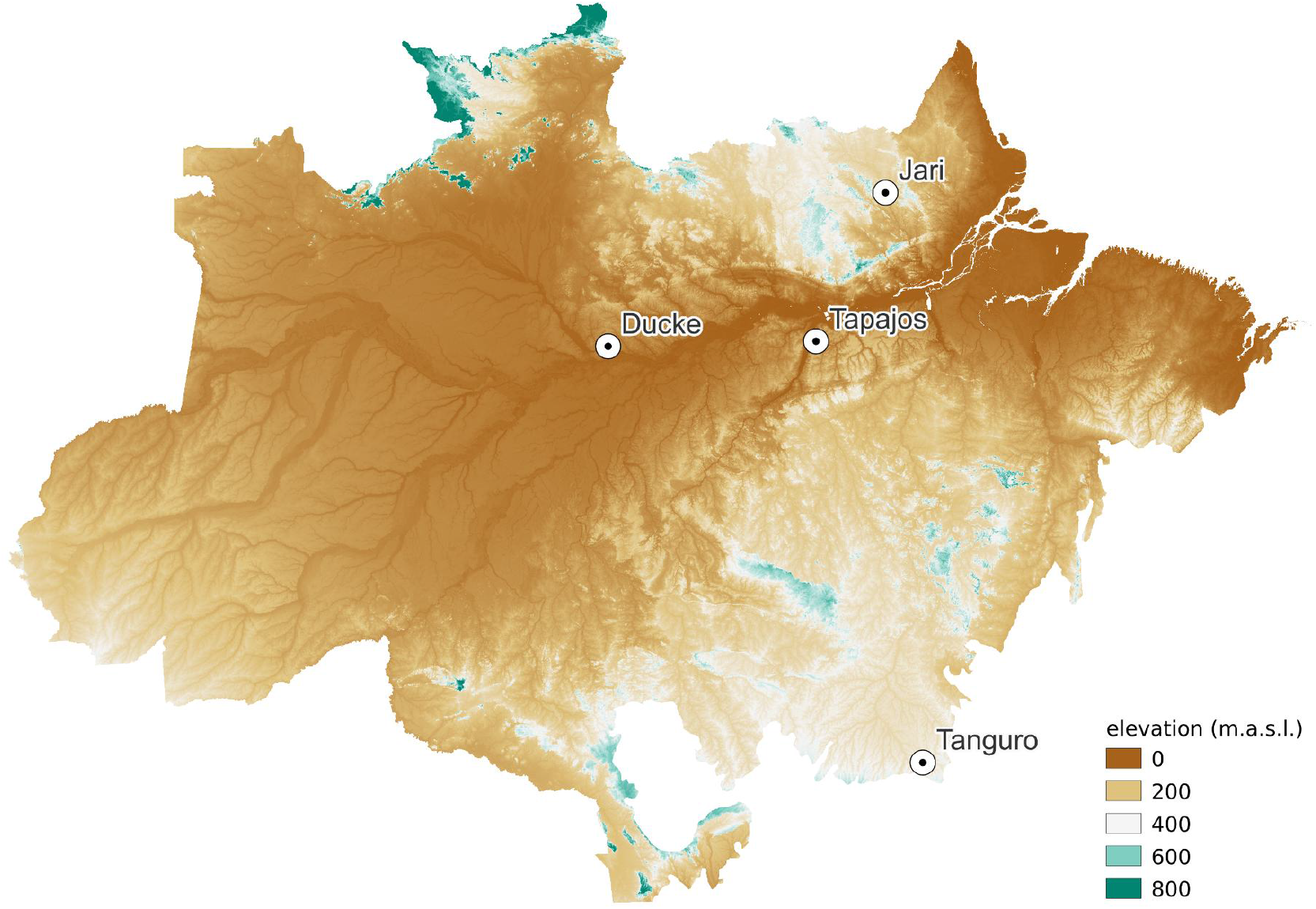
Sites distribution in the Brazilian Amazon, with terrain elevation in the background. Jari is located in the northeast, Tapajos in central-east, Tanguro in the southeast and Ducke in the central Amazon.

**Table 1.** Study sites description based on environmental layers (Gorgens et al, 2021). *Temp* indicates temperature (in °C), *season* indicates the coefficient of variation (CV in % based on monthly precipitation), *precip* indicates precipitation (in mm yr^-1^), FAPAR indicates fraction of absorbed photosynthetically active radiation (in %), v-speed and u-speed indicate northsouth and east-west average wind speeds respectively (m s^-1^), elevation indicates the ground elevation from sea level (in m), clear days indicates the number of days without clouds in a year (days yr^-1^), days > 20mm indicates number of days in a year with precipitation greater than 20 mm, and maximum height indicates the approximate maximum tree height by site.

| Site | Jari | Ducke | Tapajos | Tanguro |
| --- | --- | --- | --- | --- |
| Temp. max. (°C) | 32.0 | 32.6 | 31.5 | 34.0 |
| Average temp. (°C) | 25.4 | 27.0 | 25.1 | 25.2 |
| Temp. season. (CV%) | 6.0 | 4.9 | 5.4 | 10.2 |
| v-Speed (m s<sup>-1</sup>) | 1.4 | 2.2 | 1.8 | 3.2 |
| u-Speed (m s<sup>-1</sup>) | 0.4 | 1.7 | 1.4 | 2.1 |
| Average precip. (mm yr<sup>-1</sup>) | 2335 | 2194 | 2012 | 1618 |
| Precip. driest (mm month<sup>-1</sup>) | 60.7 | 76.0 | 45.0 | 1.0 |
| Precip. wettest (mm month<sup>-1</sup>) | 346.3 | 285.0 | 368.0 | 284.0 |
| Precip. season. (CV %) | 48.3 | 41.0 | 65.0 | 80.0 |
| Lightning rate (flash yr<sup>-1</sup>) | 6.9 | 37.0 | 11.8 | 19.6 |
| FAPAR (%) | 83.4 | 84.9 | 85.0 | 63.0 |
| Elevation (m.a.s.l.) | 262.3 | 104.0 | 198.0 | 327.0 |
| Clear days (days yr<sup>-1</sup>) | 51.3 | 68.4 | 53.2 | 167.2 |
| Days >20 mm (days yr<sup>-1</sup>) | 37.9 | 36.6 | 44.0 | 31.2 |
| Maximum height (m) | 75 | 55 | 58 | 40 |

### Airborne laser scanning data collection

The laser scanning data were acquired in two campaigns. The campaign was carried out in the years 2017 and 2018, and the second in 2020. The minimum return density was of 4 points per m^2^, which was adequate to quantify canopy height in Amazon forests (Silva et al 2017). Detailed information about the flight parameters is available in Leitold et al (2018) and Cordeiro (2021) for Sustainable Landscape dataset, and in Gorgens et al (2019) for Estimation Biomass for Amazon dataset. The digital terrain model was created for each cloud based on points classified as ground by the Cloth Simulation Function algorithm (Zhang et al 2016) implemented in lidR package (Roussel et al 2021) at a resolution of 1 m. The digital canopy height model of each flight was created from the normalized cloud, retaining the maximum height (point elevation minus ground elevation) for each 1 m grid cell.

### Detecting gaps in ALS data

The gaps were identified and their area computed from the canopy height model. Following Hunter et al. (2015), we define gaps as contiguous areas with canopy height equal or less than 10 m height and area equal to or greater than 10 m^2^. Hunter et al (2015) also studied two of the sites included in the present study (i.e. Ducke and Tapajos). Gaps were grouped into three classes: small (< 100 m^2^), medium (100 - 500 m^2^) and large (> 500 m^2^) gaps based on prior studies (Leitold et al. 2018; Chambers et al 2013). Leitold et al. (2018) compared gaps identified by lidar and the necromass found in coarse woody debris surveys at the Tapajos site to separate branch fall (≤ 100 m^2^), single treefall or multiple branch fall (100–500m^2^), and multiple treefall events (≥ 500 m^2^). For each site, the gap distribution from the two surveys were compared using a chi-squared test. We were also able compared whether the distribution at each site changed over time (i.e., between consecutive campaigns).

### Gap dynamics

To assess the gap dynamics, we identify four transitions between closed canopy (nongap) and gap states (Figure 2). The base state for an area of forest is a closed canopy. From that state, an area can move to the gap state by the formation of a new gap (1); an existing gap may expand (2) or persist (3); finally, a gap may recover to the state of close canopy (4). We computed state transitions by overlapping the gap polygon layers from the first and second campaigns. Formation quantifies new gap area formed after the first survey, that is not contiguous with a prior gap. Gap expansion occurs when the first state was closed canopy, the second state was gap, and it is contiguous with an existing gap. Persistence is defined for a polygon identified as a gap in the first flight and that stayed gap in the second flight. Recovery is defined as an area of gap in the first flight that becomes closed canopy in the second flight. Full gap recovery and partial gap recovery are not discriminated by the overlay method applied. The results were presented in area of gaps per hectare, per year (m2 ha^-1^ yr^-1^), and in number of gaps per hectare, per year (n ha^-1^ yr^-1^) to normalize the study area sizes and varying periods between consecutive surveys. Most time differences between flights were about three years. For Tanguro, the difference was about two years.

**Figure 2.**
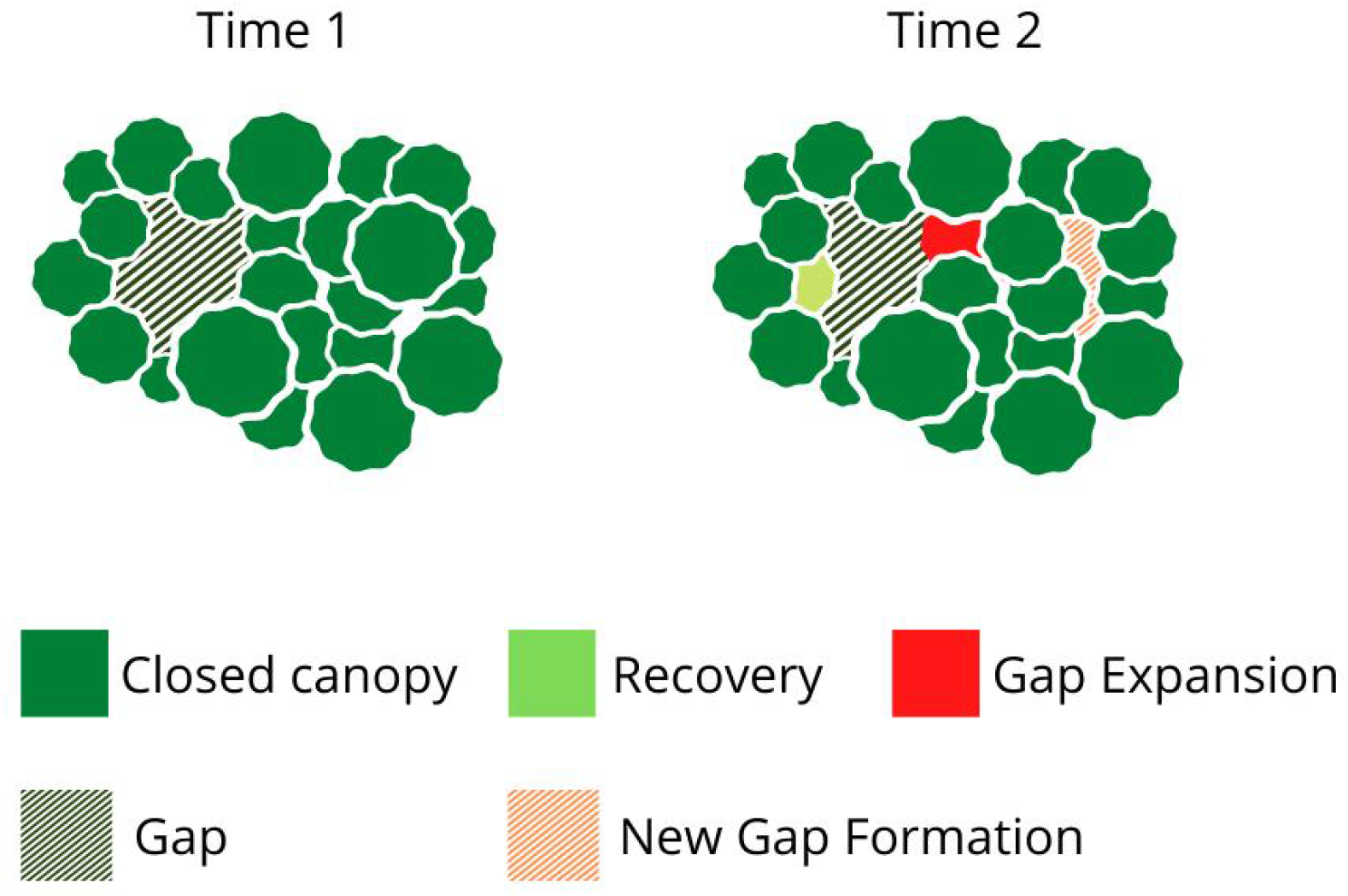
Forests are composed of two states, gap (striped) and closed canopy (solid). When closed canopy becomes gap, we call this *new gap formation.* Old gaps may increase in area *(gap expansion)* or lose area to the closed canopy by regrowth *(recovery).*

In addition to accounting for transitions, we investigated the size of new gaps (gaps not detected in the first flight) classifying them into small, medium or large. The dynamics of new gaps was expressed as area of gaps per hectare per year (m^2^ ha^-1^ yr^-1^), and as number of gaps per hectare per year (gaps ha^-1^ yr^-1^). Finally, we computed the rates of change in canopy height by subtracting the canopy height model derived from the second flight by the canopy height model derived from the first flight, and dividing them by the period between flights. The rate expresses the change in height by year (m yr^-1^). We computed the average for pixels within respective polygons of closed canopy, persisting gaps and recovered gaps. Data analysis was performed in R 4.1.1 (R Core Team, 2021) and QGIS 3.18 (QGIS Development Team, 2021).

## Results

Gap size frequency in all studied sites followed a power-law distribution with a high proportion of small gaps, and a small proportion of large gaps (Figure 3). While Ducke did not have gaps greater than 2,500 m^2^, Jari had gaps with up to ~12,500 m^2^. All the sites had a similar proportion of gaps among size classes (Q1, p-value for χ2 ~ 0.26 for the first flight and 0.18 for the second flight). Even though the forest in Jari contained larger gaps, they still have a small proportion of large gaps overall.

**Figure 3.**
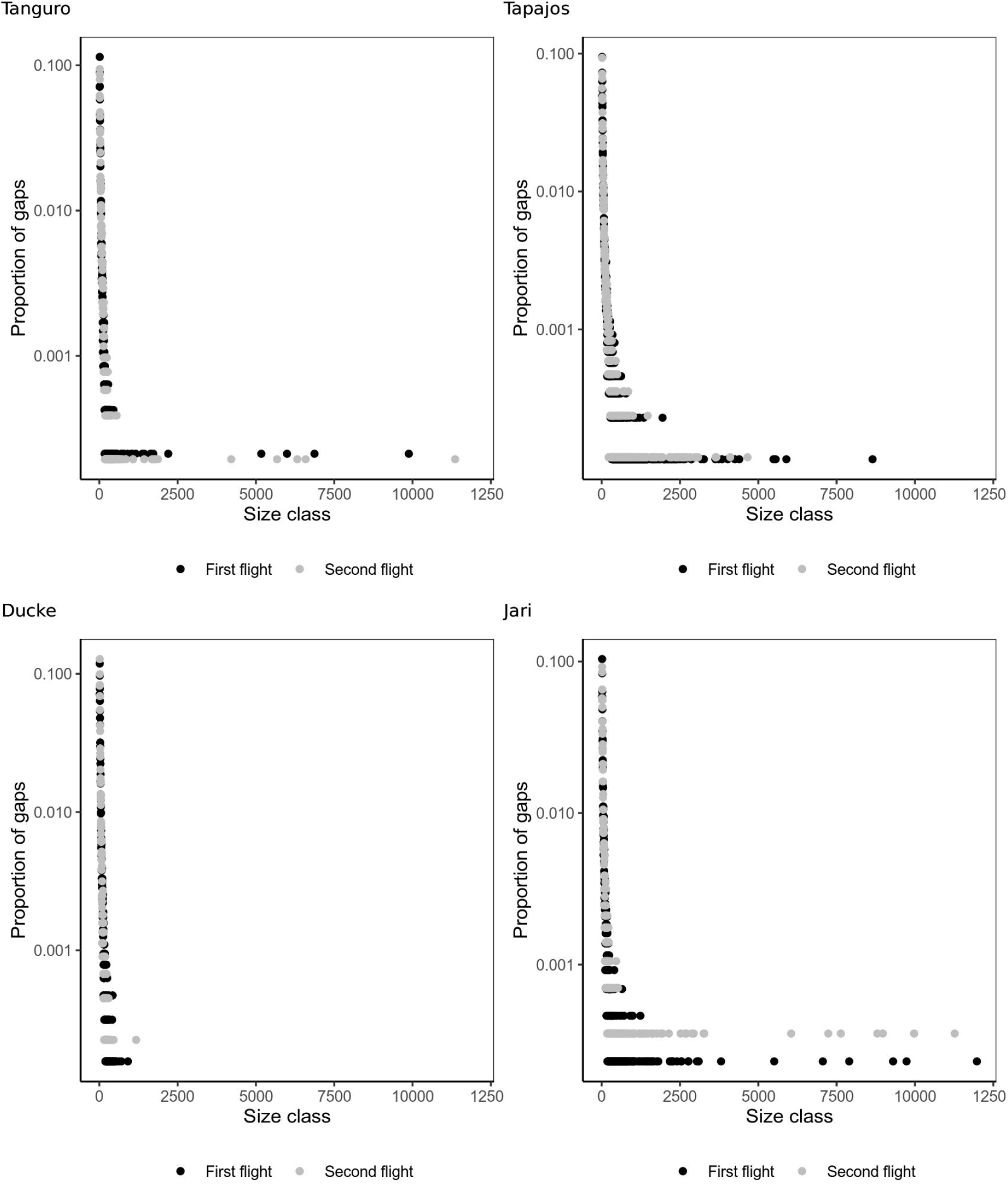
Proportion of gaps in terms of frequency as a function of the gap size per study site.

Comparing the monitored years, we found changes in the gap distributions for all sites. The changes were significant in Tapajos and Ducke (p-value for χ2 < 0.05), while Tanguro and Jari did not show statistically significant differences between the first and second flight gaps distributions (p-value for χ2 ~ 0.5).

The density of gaps observed in this study varied between 3.50 to 8.72 gaps ha^-1^. The largest number of gaps was observed in the Tapajos, for both surveys (8.52 gaps ha^-1^ in the first flight and 8.25 gaps ha^-1^ in the second flight) and the lowest was observed in the Ducke and Jari (first flight in Ducke: 5.26 gaps ha^-1^ and second flight in Jari: 3.5 gaps ha^-1^). During the monitored period, only the Tanguro site showed an increase in the number and area of gaps, which changed from 7.99 gaps ha^-1^ (area: 399 m^2^ ha^-1^) to 8.72 gaps ha^-1^ (area: 472 m^2^ ha^-1^) (Table 2).

**Table 2a.**
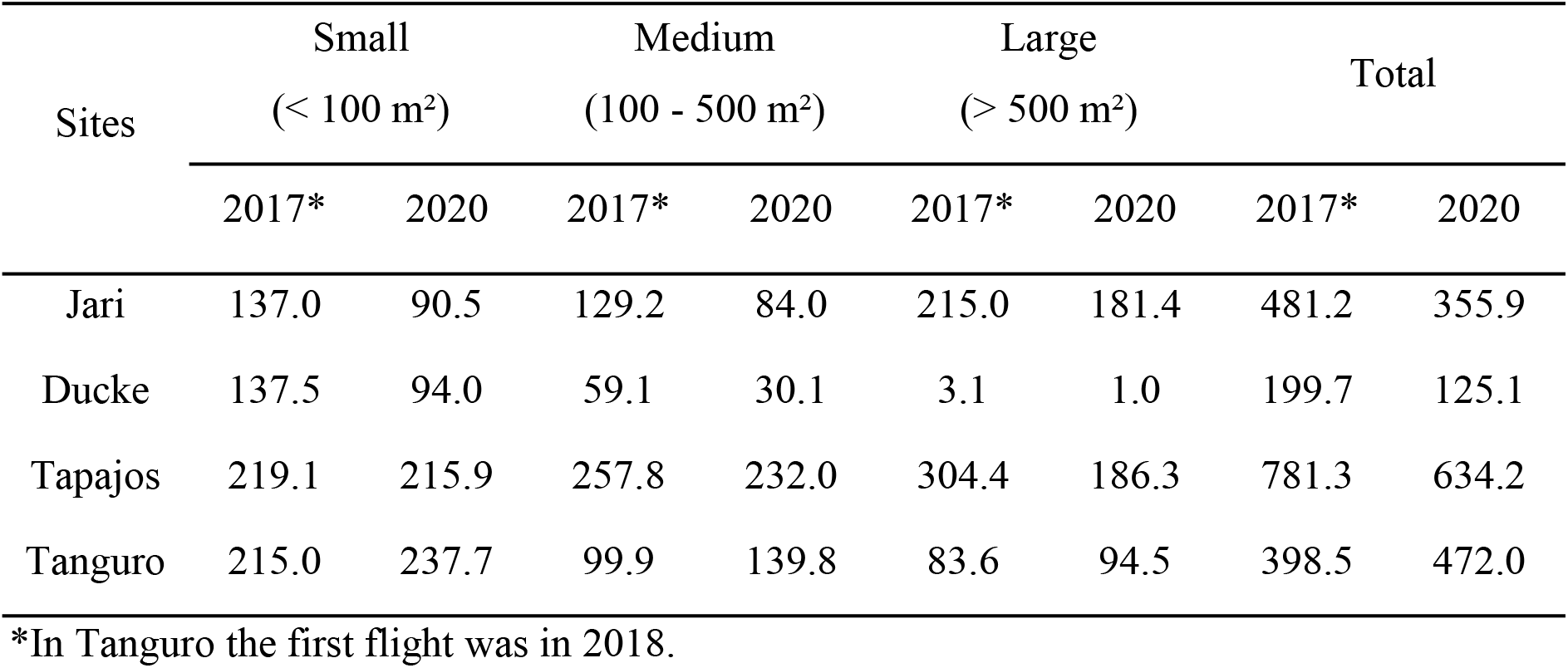
Area of gaps per hectare of forest per size class in the two monitoring flights (m^2^ ha^-1^).

**Table 2b.** Number of gaps per hectare of forest per size class in the two monitoring flights (gaps ha^-1^).

| Sites | Small<br>(< 100 m<sup>2</sup>) |  | Medium<br>(100 - 500 m<sup>2</sup>) |  | Large<br>(> 500 m<sup>2</sup>) |  | Total |  |
| --- | --- | --- | --- | --- | --- | --- | --- | --- |
|  | 2017* | 2020 | 2017* | 2020 | 2017* | 2020 | 2017* | 2020 |
| Jari | 4.57 | 3.01 | 0.65 | 0.41 | 0.13 | 0.08 | 5.35 | 3.50 |
| Ducke | 4.91 | 3.50 | 0.35 | 0.18 | 0.005 | 0.001 | 5.26 | 3.68 |
| Tapajos | 7.02 | 6.91 | 1.24 | 1.17 | 0.26 | 0.17 | 8.52 | 8.25 |
| Tanguro | 7.36 | 7.87 | 0.58 | 0.80 | 0.04 | 0.05 | 7.99 | 8.72 |

\*In Tanguro, the first flight was in 2018.

Unsurprisingly, gap number was dominated by relatively small gaps. For all sites in both surveys, the percentage of gaps <100 m^2^ ranged from 82% (Tapajos 2017) to 95% (Ducke 2020). Ducke always had the largest proportion of small gaps, and the Tapajos site always had the largest proportion of large gaps (Table 2a). The Tapajos site had the largest number of gaps on average and also, by far, the largest gap area. The distribution of gap area was relatively even across the small, medium, and large categories in Tapajos and Jari, whereas at Tanguro and especially at Ducke, medium and large gaps accounted for only a small portion of the gap area (Table 2a).

The highest new gap formation rate was observed in Tanguro with 46.52 m^2^ ha^-1^ yr^-1^, followed by Jari and Tapajos with ~33.5 m^2^ ha^-1^ yr^-1^, and Ducke 15.89 m^2^ ha^-1^ yr^-1^ (Table 3). Tapajos and Tanguro had the greatest gap expansion rates, followed by Jari and Ducke (Table 3). Considering the increase in gap area as formation plus expansion, Tanguro and Tapajos had more than 1% of total forest canopy area converted to gaps annually, with Jari at about 0.8% and Ducke well below that level at about 0.3%.

**Table 3.** Area of gap transitions per hectare per year (m^2^ ha^-1^ yr^-1^) and the transition probability (percentage of total gap area mapped in parentheses).

| Site | Formation | Expansion | Persistence | Recovery |
| --- | --- | --- | --- | --- |
| Jari | 35.84 (14.8%) | 41.04 (17.2%) | 42.18 (17.8%) | 118.88 (50.1%) |
| Ducke | 15.89 (14.8%) | 11.99 (11.2%) | 21.63 (20.1%) | 57.90 (53.9%) |
| Tapajos | 32.22 (8.6%) | 82.41 (21.9%) | 97.84 (26.0%) | 163.66 (43.5%) |
| Tanguro | 46.52 (14.6%) | 71.79 (22.5%) | 117.81 (37.0%) | 82.29 (25.8%) |

Stasis in the canopy is the rule over short time periods. For the duration of our investigation (2-3 years) at four sites, between 97.2% and 99.1% of the canopy remained in the same state that it began annually (i.e. closed canopy remained closed canopy and gap remained gap). Nevertheless, there were measurable canopy dynamics and the transition probabilities between canopy states were significantly different among regions (p-value for χ2 ~ 0.0011) (Table 3). Jari showed a proportion of gap recovery of 50.1% and Ducke 53.9%. Tapajos had a recovery rate of 43.5%, while Tanguro showed 25.8% (Table 3). Compared to the other sites, Tanguro had a high proportion of gaps persisting (37%) and recovering (22.5%). Tapajos had a persisting proportion of 26.0%, followed by Ducke (20.1%) and Jari (17.8%). For recovery, Tapajos showed a proportion of 21.9%, followed by Jari (17.2%) and Ducke (11.2%). This had 0.44 new gaps ha^-1^ yr^-1^, and Tanguro had more than 1.4 gaps ha^-1^ yr^-1^. Although Jari and Ducke showed low and similar disturbance rates in terms of number of gaps (i.e., gaps ha^-1^ yr^-1^), Jari had a considerably largest area of new gaps (13.3 m^2^ ha^-1^) (Figure 4).

**Figure 4.**
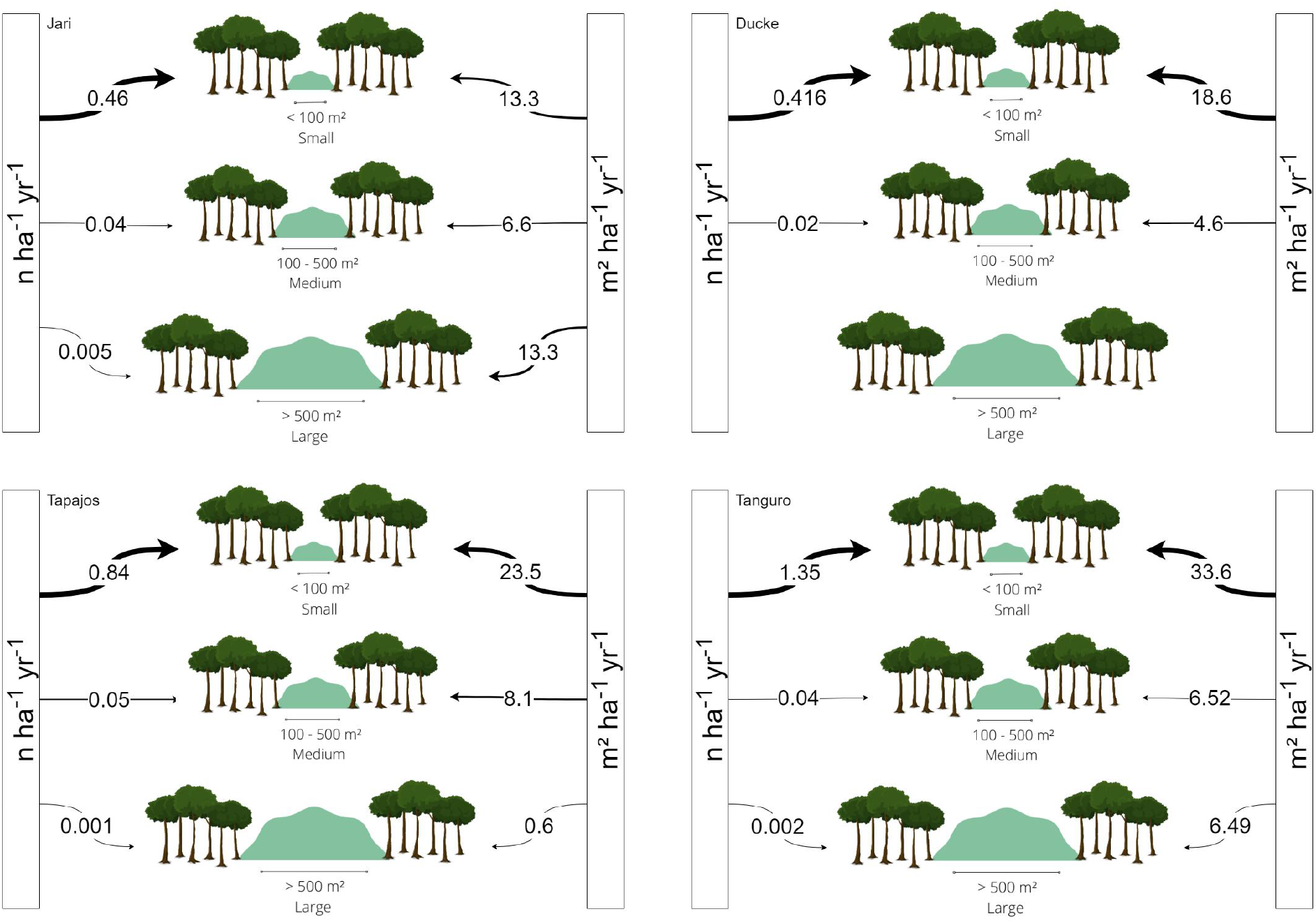
Diagram expressing area and number of new gaps (formation) per hectare per year by gap size class.

The rate of recovery exceeded the rate of new gap opening (formation plus expansion) by 43% to 108%, with the exception of Tanguro where recovery was only 70% as great as the sum of formation plus expansion. Proportionally, the Tanguro site had the greatest gap persistence (37%). The canopy height changes computed from our lidar data confirms that the fast recovery was associated with higher canopy changes. Jari exhibited the highest growth rate for closed canopy, reaching on average 0.71 m yr^-1^, followed by Ducke (0.35 m yr^-1^), Tapajos (0.15 m yr^-1^) and Tanguro (0.06 m yr^-1^) (Table 4). The higher the changes in canopy height, the smaller is the lifetime of gaps.

**Table 4.** Net canopy height changes (m yr^-1^) by gap dynamics.

| Sites | closed canopy | persisting gaps | recovered gaps |
| --- | --- | --- | --- |
| Jari | 0.71 | 0.47 | 4.46 |
| Ducke | 0.35 | 0.30 | 2.07 |
| Tapajos | 0.15 | 0.48 | 3.72 |
| Tanguro | 0.06 | 0.25 | 2.31 |

## Discussion

Canopy gaps emerge from basic properties of forest communities, such as the size, number and demography of individual trees (Jucker et al 2021). Our studied sites are covered by old-growth forests, with a low frequency of large disturbances (< 1% for all sites, Figure 4). All sites had a high proportion of small gaps likely associated with crown or branches fall. Medium size gaps are related to multiple branch fall or single treefall. Large gaps are related to the mortality of tree clusters often promoted by exogeneous disturbances such as extreme rain and wind, which cause snapping and/or uprooting (Puig 2009, Bottero et al 2011, Leitold et al 2018, Chambers et al. 2013; Marra et al. 2014; Negrón-Juarez et al. 2011).

Our studied sites showed gaps per hectare ranging between 3.5 and 8.72 gaps ha^-1^, and a rate of new gaps ranging between 0.44 and 1.4 gaps ha^-1^ yr^-1^. Tanguro had the largest gap fraction among the sites. This was the site with the lowest recovery (25.8%), and the highest persisting (37%) and recovery rates (22.5%). Tanguro is located in the southeast of the Amazon, close to the transition to the *Cerrado* biome, with the strongest climatic variation, the lowest precipitation, the highest number of sunny days, and the highest maximum temperature. Forests bordering the transition zone with the Brazilian *Cerrado* were previously reported as having higher gap dynamics than rainforests (Dalagnol et al 2021). Forests in this region may be especially vulnerable to degradation due to the synergy of human an extreme natural-disturbances such as windthrows (Silverio et al. 2019).

The static size distributions of gaps for our four sites are similar to those found in other remote sensing studies of tropical forests (Kellner and Asner 2009; Asner et al. 2013). We found no significant difference among the proportion of gaps by size classes, although large gaps were more prominent in the Tapajos and Jari sites. Fisher et al. (2008) suggested that gap frequency distributions could be used to model disturbance frequency. Reis et al. (2021) analyzed the gap size-frequency distributions across 650 lidar transects covering the Brazilian Amazon forest. They observed that human disturbed forests had a higher proportion of large gaps than intact forests. In addition, areas with strong wind gust speeds, frequent lightning and more severe water shortages had a higher proportion of large gaps.

Jucker (2022) questioned what can we learn from these gap size frequency distributions. He concluded that we need more modeling studies to understand the controls of tropical forest gaps. We add that more empirical data is required for reliable assessments and models of mechanisms of tree mortality influencing the size distribution and dynamics of canopy gaps. Gaining further understanding on the size distribution and dynamics of gaps requires sorting out the influence of branch falls, single and multiple treefalls as these disturbance and mortality modes overlap (Leitold et al. 2018). Understanding how gap size and frequency influence recovery and successional changes in structure and diversity is also crucial (Denslow et al. 1998; Chambers et al. 2013; Marra et al. 2018). Modeling studies alone will not explain gap frequency distributions without data constraints.

In order to better understand forest gap processes, we focused on the dynamics of gaps using measurements from two time periods. Multitemporal gap turnover measurements using lidar are rare in tropical forests (see Hunter et al. 2015 and Leitold et al. 2018). Over the short-time interval of our study, the studied forests were clearly not in a steady-state. At steady-state the ratio of gap *recovery* to the sum of gap *formation* plus *expansion* should be equal to one. We found that the ratio of gap recovery to the sum of formation plus expansion ranged from 1.4 to 2.1 for Tapajos, Jari, and Ducke sites. Only Tanguro had a ratio smaller than one (0.7). On the other hand, prior work by Hunter et al. (2015) for Ducke and Tapajos sites using nearly identical methods showed very similar rates of gap formation plus expansion (26.0 and 30.5 m^2^ ha^-1^ y^-1^ respectively) for the time period 2008 to 2012 compared to those found in this study. It is reasonable for recovery to dominate at most sites in this study because we expect higher than normal rates of disturbance associated with the strong El Nino related drought in 2015-2016 prior to our first survey (Leitold et al. 2018).

Where new gaps were not created, the average change in canopy height was always positive, which indicates that even in absency of detectable disturbances, old-growth forests are continuously growing. The height gains were greatest for recovered gaps (>2 m y^-1^) probably because of a contribution from lateral growth of trees at the surrounding canopy (Kellner and Asner 2014; Hunter et al. 2015). Height change in gaps has two components: lateral growth from the gap edges and vertical growth from below. In a previous study in Ducke and Tapajos, lateral growth accounted for only 6–10% of gap closure (Hunter et al 2015). Closed canopies and persisting gaps had similar slow rates of height increase (<1 m y^-1^) because those areas contained a mixture of loss and gain pixels. In general, aggrading sites where the ratio of gap recovery to the sum of formation plus expansion was greater than one (Jari, Ducke, and Tapajos), had larger annual height gains compared to the degrading site with lower than one value (Tanguro). Apart from different forest structure and species composition, the latter site is considerably drier than the others and water limitation may restrict height growth.

Based on prior studies, we know that the rate of canopy turnover and gap formation varies greatly from year to year. Observing the fall of coarse woody debris in the Tapajos forest, Palace et al. (2008) found that necromass production at one of his two 100-ha study sites was dominated by a single storm event. This result is similar to the findings of Araujo et al. (2021) in the 50-ha plot at Barro Colorado Island, Panama. They found that 20% of all canopy damage registered over 5 years occurred in a single 42-days period. Canopy damage and tree mortality are associated with episodic events such as lightning, convective storms, and severe winds (Yanoviak et al. 2020; Negron-Juarez et al. 2018; Espirito-Santo et al. 2010; Silverio et al. 2019). These results indicate that long-term monitoring is also key for achieving consistent and reliable insights.

Comparison across sites and studies is difficult because of different definitions for gaps and canopy disturbance. The definitions vary according to both horizontal (plane area) and vertical (depth within the canopy) dimensions. Plane area varies from 4 m^2^ (Leitold et al. 2018) to 25 m^2^ for studies using photogrammetry (Araujo et al. 2021; Cushman et al. 2022). Emphasis on different dimensions depends on the purpose of the study. For example, Leitold et al. (2018) focused on carbon cycling effects of canopy disturbance. They selected a minimum canopy area of 4 m^2^ and a minimum height change of 3 m in order to capture branch disturbance. Studies that focus on forest regeneration, and especially illumination conditions at the forest floor will tend to use larger horizontal dimensions and greater depth within the canopy. All other things being equal, turnover rates will be larger with smaller area and shallow depth constraints.

This study agrees with prior work showing that tropical forest gaps are contagious (Young and Hubbel, 1991; Hunter et al. 2015). The annualized area of gap expansion was larger than gap formation at three of four sites investigated in this study. Only at Ducke, where the canopy was least dynamic, gap formation exceeded the gap expansion area. Hunter et al. (2015) suggested that edge effects may explain gap contagiousness. More recently, Arellano et al. (2019) discovered another process that could lead to gap contagiousness. They demonstrated that crown damage was a strong predictor of tree mortality. Tree-fall or branchfall events can easily damage neighboring trees making them more likely to die, subsequently increasing contagious canopy turnover.

Gap formation and canopy turnover measured by lidar are linked to tree mortality but they are not identical. Branch fall or more severe canopy damage is recorded by lidar but is not accounted for as mortality in forest inventory surveys. Often, these events are also not quantified in biomass/carbon inventories (Peixoto 2021). In contrast, tree mortality will not necessarily be registered in lidar surveys of forest turnover. Trees that die standing may not immediately create a gap in lidar surveys and may never lead to gap opening if regrowth around the dead tree and decomposition of the snag follow similar rates. Nevertheless, recent studies found that field based mortality was related to lidar measures of canopy turnover across the Brazilian Amazon (Dalagnol et al. 2021) and in French Guiana (Huertas et al. 2022). We believe that relations between the two quantities (formation and turnover) may be improved by considering regional variations in the modes of tree death (Esquivel-Muelbert et al. 2020).

## Conclusions

Our sequence of two airborne laser scanning surveys allowed us to expand the investigation about gap-fraction, quantifying the gap dynamics composed by four stages: formation, expansion, persistence and recovery. We found that all of our sites were far from steady-state over the 2 to 3-year interval of the study. Out of four, forest recovery area exceeded the area of new gap formation in three sites. Where gaps formed, there was a tendency for them to be located adjacent to existing gaps (gap expansion was more common than new gap creation) suggesting that gap opening is a contagious process.

Gaps are the manifestation of how disturbances disrupt forest landscapes, opening the canopy to sunlight and trigging succession, which increase heterogeneity, diversity and complexity to forest canopies. The concept of stability reflects the tendency of a system to quickly return to a position of equilibrium when disturbed. We show that gap dynamics varied among sites, with one example of a low recovery rate contrasted to sites with faster recovery. Looking only at the distribution of gaps produce an incomplete picture of the forest dynamics. Both gap formation and recovery must be considered. Future studies using airborne lidar and photogrammetry at more sites over longer periods have the potential to greatly improve our understanding of forest canopy dynamics.

## Acknowledge

Funding was provided by the Coordenação de Aperfeiçoamento de Pessoal de Nível Superior Brasil (CAPES; Finance Code 001); Conselho Nacional de Desenvolvimento Científico e Tecnológico (Processes 403297/2016-8 and 301661/2019-7); Amazon Fund (grant 14.2.0929.1); National Academy of Sciences and US Agency for International Development (grant AID-OAA-A-11–00012); Universidade Federal dos Vales do Jequitinhonha e Mucuri (UFVJM); and Instituto Nacional de Pesquisas Espaciais (INPE). D. Almeida was supported by the São Paulo Research Foundation (#2018/21338-3 and #2019/14697-0). T. Jackson and D. Coomes were supported by the UK Natural Environment Research Council grant NE/ S010750/1.

## Notes

https://github.com/Gorgens/gap-dynamics

### Competing Interest Statement

The authors have declared no competing interest.

